# Noncoding elements within *MYCN* mRNA are autonomous drivers of oncogenesis in neuroblastoma

**DOI:** 10.1101/2025.01.10.632492

**Authors:** Vishwa Patel, Lorraine-Rana E. Benhamou, John T. Powers

**Affiliations:** Division of Pharmacology and Toxicology, College of Pharmacy, The University of Texas at Austin; Department of Pediatrics, Dell Medical School at The University of Texas at Austin

## Abstract

Neuroblastoma (NB) is a highly metastatic pediatric cancer arising from the neural crest lineage. Genetic amplification of the *MYCN* proto-oncogene is a defining feature of NB, present in about 20% of all cases. The *let-7* tumor suppressor microRNA targets the 3’ UTR of *MYCN* mRNA. We previously demonstrated that the 3’ UTR of *MYCN* mRNA acquires the ability to sequester *let-7* in *MYCN*-amplified (*MA*) disease, thus inhibiting its function. This work established that a noncoding element within an oncogenic mRNA can contribute independently to disease pathology and genetic patterning. To further investigate the roles of noncoding RNA elements within the *MYCN* mRNA, we engineered cells expressing either *MYCN-ORF* (MYCN open reading frame only), *MYCN-GL* (full-length MYCN mRNA from the intact genetic locus), and *Null-GL* (full-length *MYCN* mRNA variant where EGFP replaces MYCN protein). We observe that all constructs enhance growth compared to controls *in vitro* and *in vivo. MYCN-GL*-expressing cells displayed the most robust growth *in vitro* despite containing multiple regulatory RNA elements. Remarkably, the *Null-GL* construct induces cells to grow as fast or faster than *MYCN-ORF*-expressing cells. Animal studies further confirmed these observations, where the *Null-GL*-driven tumors had the highest incidence and lowest latency, followed by *MYCN-GL* and then *MYCN-ORF*. Further, through NGS analysis, *let-7, miR-101*, and *miR-34a* targets are enriched in both *MYCN-GL* and *Null-GL* expressing cells. Thus, the 3’UTR of *MYCN*, which is also targeted by these microRNAs, may interact with them in *MYCN-GL* and *Null-GL* cells to deliver a protective effect for the mRNA targets of these microRNAs. We also observed more dynamic differential gene expression in the -GL constructs than in ORF and GFP-expressing cells. In addition, *MYCN-GL* and *Null-GL* expressing cells are similarly enriched in gene ontology pathways for cancer, RNA metabolism, and microRNA processing pathways as compared to *ORF* and *GFP*. Whole genome sequencing also revealed more similarities in copy number variation in *MYCN-GL* and *Null-GL* than in *ORF* and *GFP*, suggesting that these constructs may provide selective pressure to favor specific CNV patterns. These observations show that full-length *MYCN* mRNA containing noncoding regulatory elements are more robust drivers of cell growth and oncogenicity than MYCN protein alone and provide insights into the mechanisms of oncogenic contribution. These results open an exciting door for our understanding of NB pathology and genetic patterning and have broad implications for other oncogene-driven cancers.

## Introduction

Neuroblastoma (NB) is a highly metastatic pediatric cancer derived from immature nerve cells called neural crest lineage and is most common in infants and children under five years old. The neural crest lineage usually differentiates sympathetic ganglia, chromaffin cells of the adrenal medulla, and paraganglia.^1^ It accounts for more than 7% of childhood malignancies and around 15% of cancer-related deaths in childhood. NB has less genetic disposition than adult cancers; out of known genetic disparities in NB, *MYCN* gene amplification is the most common; in fact, *MYCN* was first discovered in NB cancer.^2,3^ *MYCN* is a proto-oncogene located on chromosome 2p24.3. Commonly, *MYCN* engages in cell proliferation, differentiation, and apoptosis. *MYCN* belongs to the MYC transcription factor family. The MYC oncogene family members are deregulated in over half of all human malignancies. Genetic amplification of *MYCN* is the hallmark of NB and occurs in 20% of patients, resulting in advanced disease stages, rapid tumor progression, and poor prognosis^4^. *MYCN* amplified NB is a devastating disease with less than a 50% survival rate. *MYCN* and *c-MYC* have highly overlapping functions and are each associated with multiple human cancers^5^. Consistent with functional redundancy, *MYC* is amplified in 10% of non-*MYCN* amplified high-risk NB, driving similar outcomes. NB is, therefore, an excellent system for studying *MYCN*, especially in the context of MYCN amplification and nonamplification, and it provides potential insight into MYC family function ^6–8^.

*MYCN* is the master oncogene of NB, where its genetic amplification results in very high mRNA expression and poor patient survival^9^. *MYCN* amplification in NB patients has both MYCN protein-cording RNA and non-coding 5’ and 3’ UTRs and two introns. Despite having other components of genes, most of the proto-oncogenic studies have focused on only the protein-coding region of the genome^8,10^. This is also true for NB; most in vitro studies including two existing mouse models (TH-MYCN and Dbh-MYCN transgenic mouse model), are driven by overexpression of *MYCN ORF*^11,12^. MYCN protein-focused studies have allowed us to make great strides in understanding the normal and oncogenic functions of MYCN protein. However, neuroblastoma is a disease of few protein-altering mutations, relying instead on frequent chromosome abnormalities, such as *MYCN* amplification and periodic chromosome arm gains or losses. Many of these large-scale genetic events are categorized into distinct patient subsets. For instance, loss of chromosome arms 11q and 3p occurs in up to 60% of *MYCN* non-amplified cases and is associated with worse outcomes^13,14^. In contrast, 11q and 3p loss are mutually exclusive with *MYCN* amplified. Further, The most common chromosomal gain of 17q occurs at high frequency in all subtypes of advanced-stage neuroblastomas and has been associated with a poor clinical outcome. Also, chromosomal alterations involving gains and losses have been observed in both above-listed neuroblastoma mouse models^11^. Considering the sizeable chromosomal imbalance in NB, these abnormalities contribute to poor prognosis and disease pathogenesis.

Since the identified most common gene in NB directly connects with chromosomal imbalances, the discovery of tumor suppressor genes may be made more difficult by the likelihood that they transmit a tumorigenic effect through haploinsufficiency^15,16^. It is also likely that the sequences that cause neuroblastoma cancer are non-translated RNA sequences with significant regulatory roles rather than typical protein-coding gene sequences. About 60% of the variation in protein expression was explained by post-transcriptional mechanisms and is vital for understanding complete proteomics. The 3’ and 5’ untranslated regions (UTRs) are the domain of mRNA, control the post-transcriptional gene regulation, and engage in pre-mRNA stability and translation initiation. Moreover, multiple non-translated RNA sequences, including microRNAs (miRNA), are mapped to the previously mentioned genomic imbalance regions that play a vital role in the differentiation of the cell, and dysregulation of these sequences can have tumor suppressor or oncogenic activity in cancer^16,17^. At the same time, we need to learn the contribution of *MYCN* noncoding RNA elements to NB oncogenesis, which is entirely unknown^1819,20^.

In neuroblastoma, let-7 microRNA plays a crucial role in tumor suppression^21^. The 3’ UTR of *MYCN* mRNA is targeted by the let-7 microRNA family, which consists of nine let-7 family members (let-7a through let-7i) spread across eight separate genetic loci. Our previous work demonstrated that inhibition of let 7 is central to the emergence of NB. Also, it established that the extraordinary levels of *MYCN* mRNA in MYCN-amplified disease turn the tables on let-7, where the 3’ UTR of *MYCN* mRNA displays an autonomous on-coding function, sequestering let-7 to inhibit its function. Research in our laboratory was the first to establish an oncogenic function for a noncoding element of *MYCN* mRNA. This discovery further explained the longstanding observation of mutual exclusivity between chromosome 11q and 3p loss, and *MYCN* amplified NB, chromosomes 11q and 3p harbor two of the most highly expressed let-7 family members. *MYCN* mRNA, through its sequestration of let-7 in amplified NB, relieves the selective genetic pressure to lose these genetic loci because the function of their gene products (let-7a and let-7g) is already compromised. Moreover, our laboratory reported that *MYCN* amplification and 11q or 3p loss account for let-7 mitigation in almost 80% of NB patients^22,23^. This work established for the first time that noncoding elements of an oncogene mRNA can independently contribute to oncogenesis and genetic patterning in cancer. It further explained why a tumor suppressor protein gene was never found to explain the loss pattern of chromosome arms 11q or 3p9,12 ^23–25^.

Additional noncoding elements exist within *MYCN* mRNA aside from the 3’ UTR. Suppose the 3’ UTR alone can have the oncogenic capacity to mitigate let-7 activity and drive impactful genetic patterns within NB^26^. The effect of whole genetic-locus derived *MYCN* mRNA molecules remains unknown. Despite the growing recognition of the importance of noncoding RNAs in cancer biology, their specific roles in NB, particularly in the context of *MYCN* amplification, have been overlooked. It is conceivable that *MYCN* noncoding RNAs profoundly affect NB development and progression, either synergistically or independently of MYCN protein^27^. Exploring the landscape of *MYCN* noncoding RNA elements holds immense promise for uncovering novel mechanisms underlying NB oncogenesis and identifying potential therapeutic targets. Integrating approaches from genomics, transcriptomics, and functional studies will be instrumental in deciphering the intricate networks orchestrated by *MYCN* noncoding RNAs in NB cells. It is therefore critical to ask these questions now to investigate all the elements together, taken together, as a fully intact mRNA and as deconstructed protein coding-only and noncoding elements-only transgenes^15^. Our recent work expanded on our earlier discovery, using three genetic constructs that address the above questions. We propose to build upon this discovery and improve genetic *in vivo* modeling of pediatric neuroblastoma (NB) by expressing *MYCN* genetic-locus-based transgenes that compare and combine *MYCN* protein function with the noncoding mRNA elements present in all human NB patients.

## Results

### Unveiling the Role of Noncoding Elements in MYCN mRNA through In Vitro Studies

Understanding the disease spectrum of cancer is critically important, as its pathology is largely based on the regulatory breakdown of the central dogma. All proteins are produced through tightly controlled processes, including the accurate reading of mRNA sequences, essentially copies of the DNA-based genetic code. Any regulation loss within this pathway, beyond mere protein function, can contribute to oncogenesis. Several elements inherent to all endogenous mRNAs are key players in this process, including the 5’ and 3’ untranslated regions and introns that span the distance between a variable number of exons for any given gene. In contrast to the ORF (protein) only approach so commonly used in oncogene overexpression (***Figure 1a)***, full-length primary (1°) mRNAs play a significant role in cancer modeling. They engage in a wide range of interactions with cellular machinery, from splicing machinery to ribosomes and translation structures and the apparatus of microRNA regulation (***Figure 1b***). Almost all of those are studied individually but not to gather, which is missing out on the interaction of all those components present in the patients, so to study that interaction, We developed three *in vitro* contracts to examine the relative effects of MYCN protein and the associated noncoding elements of the endogenous *MYCN* mRNA (***Figure 1c***). They include an *EGFP* expressing control vector, a *MYCN*-ORF vector, and a *MYCN*-Genetic Locus (*GL*)vector containing all genetic elements of the endogenous human *MYCN* primary mRNA constructs, are all in the Piggy Bac (PB) transposase system, which, when transfected with a transient PB transposase expressing vector, results in highly efficient integration of PB system vectors into mammalian genomes. The transgenes are all driven by the powerful EF1α promoter and are identical except for the specific transgene payload.

**Figure 1.**
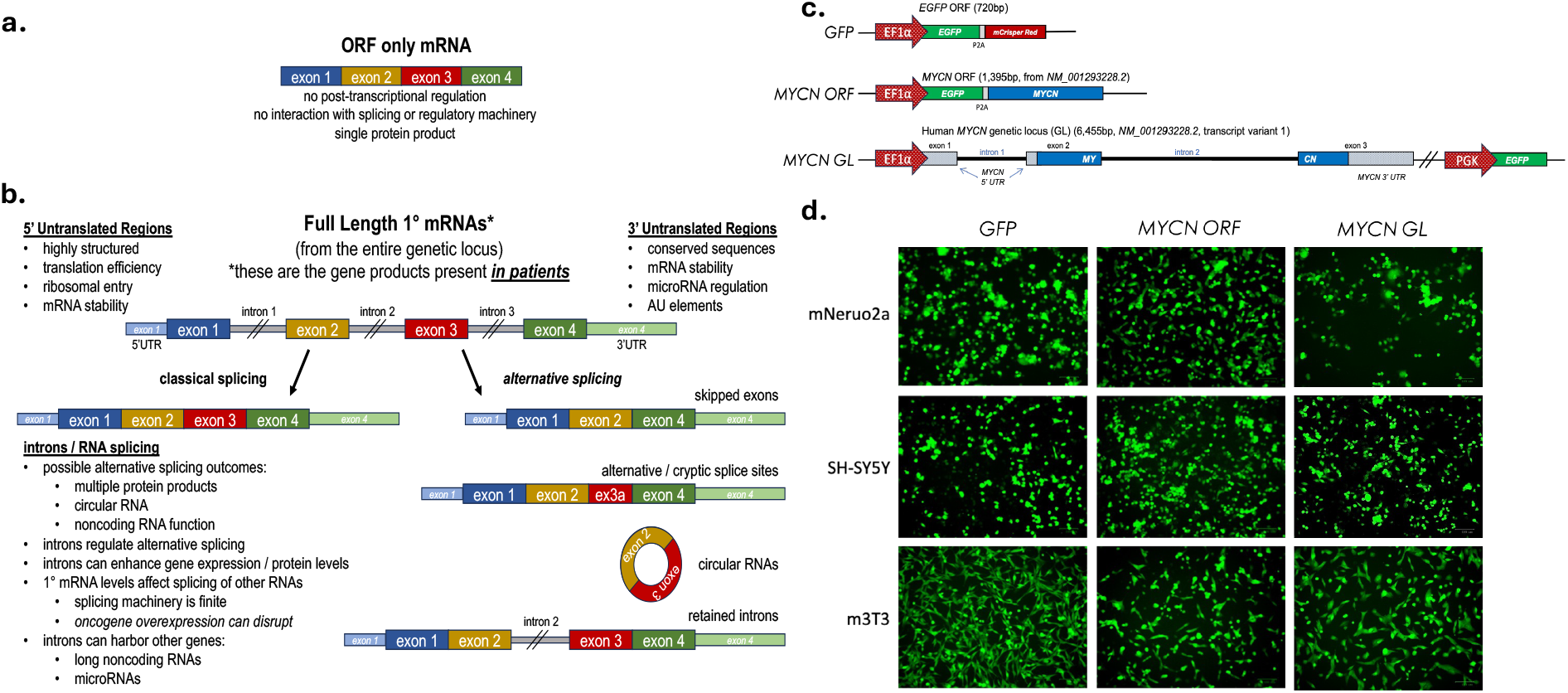
Comparison of ORF-only mRNAs with full-length, genetic locus-derived mRNAs present in patients. **a. ORF-only** derived mRNA common to many existing oncogene overexpression genetic models. **b**. Intact genetic loci yield long and complex primary mRNAs that interact with a remarkable array of processing and regulatory machinery. These genetically complete RNAs are both influenced by and apply influence back upon it. **c**. Schematic representation of MYCN Genetic constructs. All constructs contain a strong EF1a promoter in Piggy Bac transposase system vectors. The EGFP was connected to *MYCN* ORF with a P2A linker. In contrast, it was associated with the PGK promoter in *MYCN* GL **d**. Representative image of N2a, 3T3, and SHSY-5Y cells expressing *GFP, M-ORF and M-GL*,

BALB/c 3T3 (3T3) murine fibroblasts, Neuro-2a (N2a) murine, and SHSY-5Y human neuroblastoma cells were stably transfected with these constructs. 3T3s are a widely used and classical model for oncogenic transformation14, Nuero-2a is a neural crest-derived, neuroplastic, *MYCN* Non-Amplified (M-Non) murine cell line from Strain A mice widely used in neuronal differentiation, signaling pathways, and cancer studies. All three cell lines were expressing *GFP*, which indicates the expression of respective constructs in those cell lines (**Figure 1d)**. It is also an indication of well-splicing and processing by the cells. When these transduced cells were grown in 10% and 2.5% serum to observe cell growth differences. We observed significantly higher growth in *MYCN*-containing cells than GFP in all these cell types (***Figure 2a***). Interestingly, the MYCN-GL expressing cells grew faster in three cell types in both the serum conditions despite having the *MYCN* 3’UTR targeted by multiple microRNAs, including *let-7*, compared to *MYCN ORF* (***Figure 2a)***. We have investigated the expression of *MYCN* RNA in all the cell types and examined protein expression associated with the given constraint. In all three cell lines with the *MYCN ORF* that only expressed cording sequence exposed the highest amount of the MYCN protein followed by *MYCN GL* ***(Figure 2c)***, and similarly, RNA levels are similar in all three cell lines, in fact, higher in MYCN ORF ***(Figure 2d)***, which is once again aligned with the Western blot results. Also, to measure the splicing efficiency of *MYCN GL*, we have designed a specific prime set (MYCN Exon) that can target *MYCN GL* splicing efficiency, and the result shows that all the three N2a, 3T3, and SHSY-5Y cells are well-spliced by cells. These results show that MYCN mRNAs can independently provide pro-growth and oncogenic signaling, even outperforming MYCN protein.

**Figure 2:**
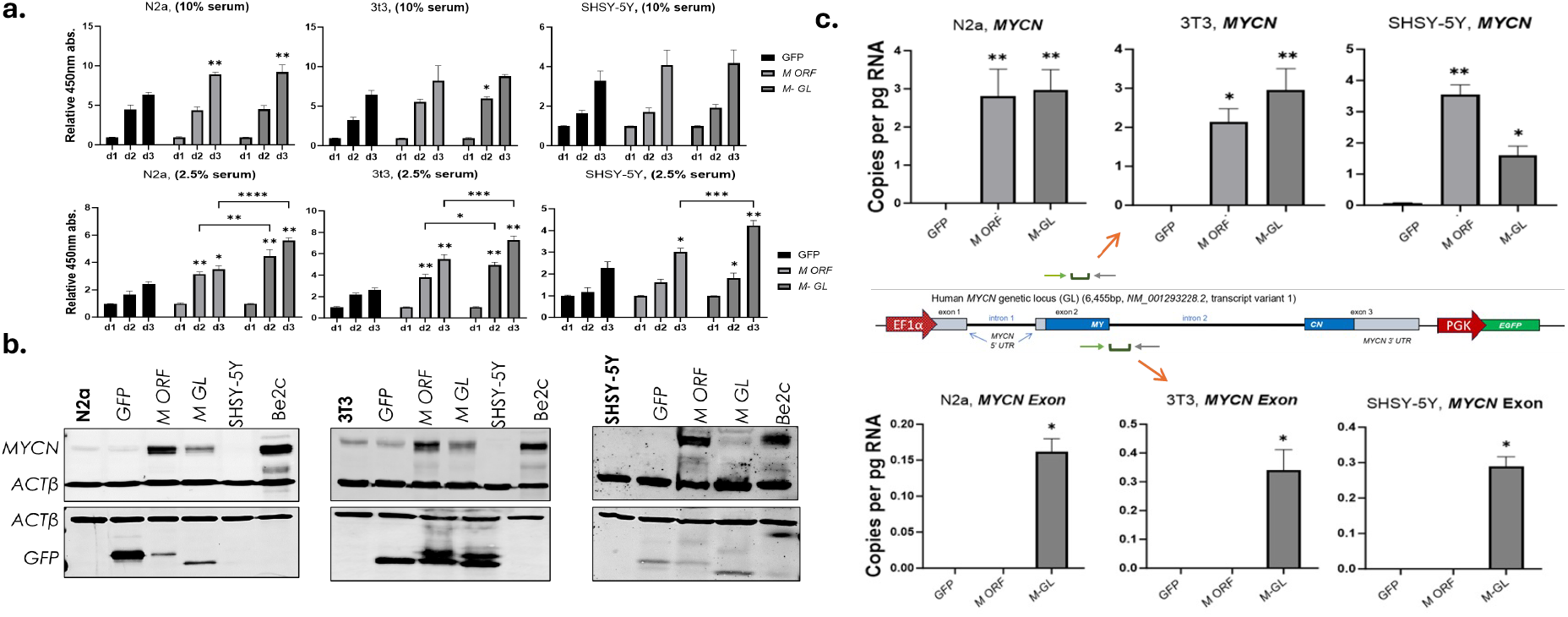
*MYCN* mRNA provides a growth advantage *in vitro*. **a**. N2a, 3T3, and SHSY-5Y cells were plated on day 0 in RPMI with 10% and 2.5% heat and inactivated FBS. Colorimetric assays for cell growth were performed on days 1, 2, and 3 (d1, d2, d3). **a**. Bars represent Mean ± SEM; significance was determined by two-way ANOVA followed by Tukey’s multiple comparison tests; *p<0.05, **p<0.01. **b**. *MYCN* ORF and genetic locus-derived full-length protein expression. N2a, 3T3, and SHSY-5Y in vitro Western blot analysis of MYCN and GFP expression in protein extracts from MYCN constructs transfected cell line. B actin was used as a loading control. **c**. *MYCN* ORF and genetic locus RNA expression levels analysis by RT-qPCR. N2a, 3T3, and SHSY-5Y cells expressing the MYCN constructs were collected to measure mRNA. Each bar represents the Mean ± SEM of copies per picogram RNA.

### Impact of Non-Coding Elements: Null GL variant of MYCN mRNA

After seeing this profound result, we have decided to make another variant of *MYCN GL*, which expresses a non-coding part of *MYCN mRNA*, which we called *Null-GL*, that is like the *MYCN GL*, but where the coding sequence of MYCN protein has been replaced with *GFP* ***(Figure 3a)*** Further, All three cell lines were expressing *Null GL* has green fluorescence, which indicates the expression of respective constructs in those cell lines and indication of well-splicing and processing by the cells (***Figure 3b*)**. Before we performed any oncogenic study, we confirmed the RNA and protein expression of the cells expressing *Null GL*. In the western blot, the *Null GL* construct does not express any MYCN protein compared to *MYCN ORF* (***Figure 3c)***. Also, the expression level of MYCN RNA in *Null-GL* is lower/Non than that of *MYCN-ORF* ***(Figure 3d)***, which once again confirms that the *Null-GL* variant of MYCN Contrast has no coding sequences. After that, we asked whether cells transduced with Null GL had any growth advantage compared to *MYCN ORF*. N2a, 3T3, and SHSY-5Y cells expressing *MYCN ORF* and Null *GL* were grown in 10% and 2.5% serum to observe cell growth differences. We observed significantly higher growth or similar growth in *Null GL*-containing cells than *MYCN ORF* in all these cell types (***Figure 3e***). Interestingly, the *Null GL* only expresses noncoding elements of MYCN mRNA, indicating that noncoding elements such as UTRs and introns also influence cancer development. The result shows that noncoding elements are equally important to study.

**Figure 3:**
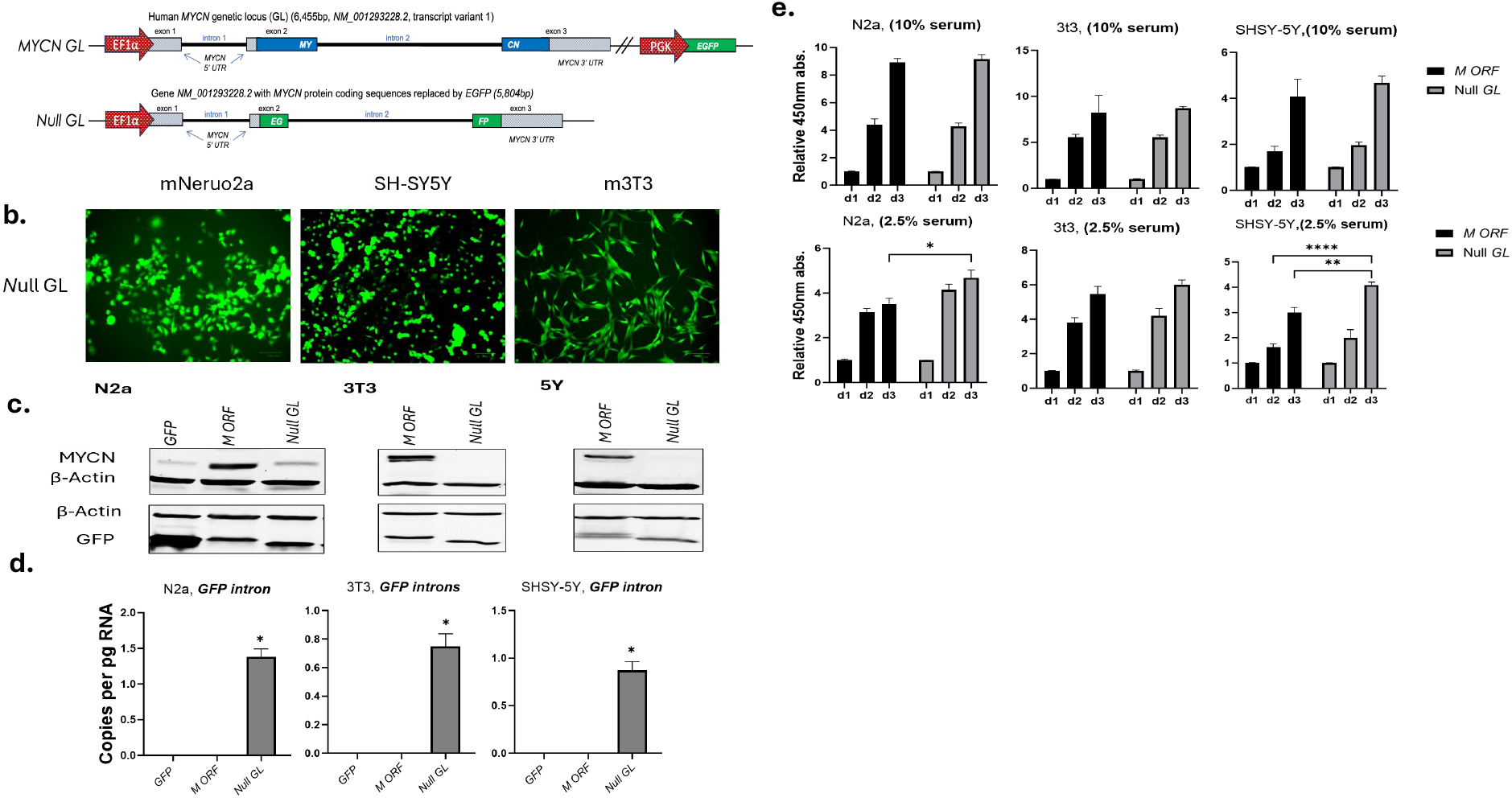
MYCN non-cording elements provide similar effects to MYCN ORF: ***a***. Schematic representation of MYCN non-cording construct, The *null-GL* was engineered to express *EGFP* instead of *MYCN ORF* in an otherwise identical locus to the *MYCN* GL construct. **b**. Representative image of N2a, 3T3, and SHSY-5Y cells expressing *null GL*. ***c***. Western blot analysis of MYCN ORF and null GL in N2a, 3T3, and SHSY-5Y cells. **d**. *MYCN null GL* RNA expression levels analysis by RT-qPCR in N2a, 3T3, and SHSY-5Y cells. Each bar represents the Mean ± SEM of copies per picogram RNA. ***e***. *No*ncoding elements of *MYCN* mRNA provide a growth advantage *in vitro* a. N2a, 3T3, and SHSY-5Y cells were plated on day 0 in RPMI with 10 % and 2.5% heat and inactivated FBS. Colorimetric assays for cell growth were performed on days 1, 2, and 3 (d1, d2, d3). Bars represent average +/-SEM; significance was represented as *p<0.05, **p<0.01 compared to control and between two groups.

### Effects of MYCN mRNA and Including Non-Cording Elements on Tumorigenesis

Once we found that ground-breaking in-vitro results. Next, we injected these transgenic 3T3 and Neuro-2a cells into syngeneic BALB/c and strain A animals, respectively. In both cases, GFP expressing cells yielded no tumors, as expected. *MYCN* ORF-expressing cells grew tumors in five out of ten injections (50%) for Neuro-2a cells ***(Figure 4a)***. Once again, the MYCN-GL-expressing cells produced more tumors in seven (70%) Neuro-2a injections in strain A Mice, and similarly, MYCN ORF-expressing cells grew two tumors out of five injections (40%) and three tumors (60%) in MYCN GL groups for 3T3 cells ***(Figure 4a)***. Also, as shown in ***(Figure 4b, c)***, tumor incidence and Volume were higher in *MYCN GL* and *Null GL* than in *MYCN ORF*. This confirms our in vitro growth pattern and further suggests that full-length *MYCN* mRNA expression from the entire genetic locus is a more robust driver of growth and oncogenicity than protein-only expression. Further, more surprisingly, we have seen *Null-GL* expressing cells that grew tumors in eight out of ten injections for N2a and three out of five injections for 3T3 cells ***(Figure 4a)***. This additional result also emphasized that the non-coding element of MYCN mRNA is equally powerful in causing cancer. We are currently identifying the Molecular mechanisms associated with MYCN mRNA in NB by analyzing the Next-generation sequencing (NGS) and Copy number variation (WGS) analysis of cells expressing this construct. This will allow us to study miRNA enrichment association and differential splicing.

**Figure 4.**
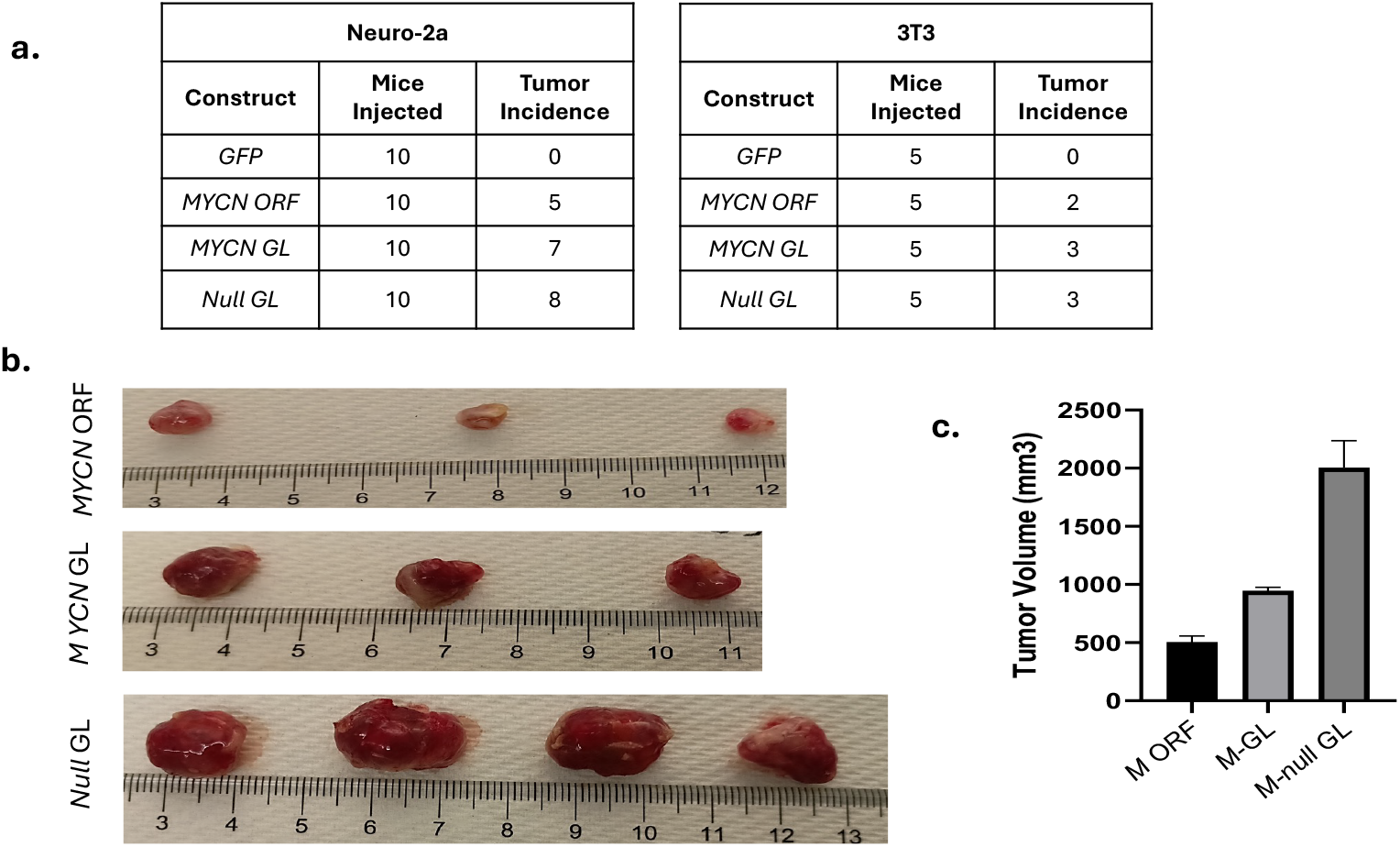
Tumorigenic effects of MYCN mRNA and their elements: ***a***. One (N2a) and two (3T3) million cells expressing the indicated constructs were injected subcutaneously into WT, *immunocompetent* Strain A, and Balb/c mice, respectively. The table represents the number of mice injected in each group, and tumor incidence was determined in 10-14 days. **b**. Presentation of tumor image from strain A mice were injected with N2a *MYCN ORF, MYCN GL*, and *Null GL* cells **c**. The bar graph represents the Difference in tumor volume (mm3) in N2a MYCN ORF., MYCN GL, and Null GL groups.

## Discussion

Neuroblastoma (NB) stands out among childhood malignancies due to its complex genetic underpinnings, with the amplification of the MYCN gene serving as a pivotal factor in disease progression. MYCN, a proto-oncogene located on chromosome 2p24.3, plays a multifaceted role in cellular processes such as proliferation, differentiation, and apoptosis. Its amplification is observed in approximately 20% of NB cases. While MYCN’s protein-coding role has garnered significant attention, recent research underscores the critical involvement of noncoding elements within MYCN mRNA in driving NB oncogenesis. These noncoding regions, including the 3’ and 5’ untranslated regions (UTRs) and introns, exert regulatory control over gene expression and post-transcriptional processes. Moreover, noncoding RNAs such as microRNAs (miRNAs) mapped to genomic imbalance regions contribute to tumor suppressor or oncogenic activity in NB ***(Figure1b)***. The discovery of MYCN’s noncoding elements as independent drivers of oncogenesis sheds new light on the complexity of NB pathophysiology and underscores the importance of holistic approaches to cancer research.

MYCN amplification (M-Amp) in NB often has up to hundreds of copies of MYCN, which results in extremely high levels of MYCN mRNA expression. Interestingly, M-Amp disease has relatively few chromosomal abnormalities compared to MYCN Non-amplified disease, which has comparatively many. Further, existing models of MYCN-driven murine neuroblastoma express MYCN ORF-only transgenes ***(Figure 1a)***. In addition, this model lacks UTRs of mRNAs that regulate mRNA stability, translation efficiency, and localization. While 5’ UTRs influence ribosomal entry and translation initiation, 3’ UTRs modulate mRNA stability via microRNA regulation. Also, these models broadly downregulate *let-7* and/or genetically lose *let-7* in patterns like non-amplified human tumors, which calls into question whether these mice faithfully model MYCN amplified disease^29,30^ Neglecting these elements in ORF-only models hampers understanding of cancer biology, especially regarding microRNA-mediated oncogene regulation. Integrating full-length mRNA expression in genetic models offers a more accurate representation, enhancing translatability and aiding therapeutic development. The gap between pre-clinical and clinical trial outcomes underscores the need for improved translational research methods. Incorporating mRNA regulatory elements in genetic models provides a more faithful representation of disease biology, facilitating better comprehension and prediction of therapeutic responses.

Our experimental findings from in vitro results suggest that MYCN GL has all the comportment of MYCN mRNA and has more cell growth advantage than MYCN ORF-only cells, encompassing protein sequence ***(Figure 2)***. In vivo models further support the oncogenic potential of MYCN mRNA in tumor development ***(Figure 4)*** in NB. Transgenic cell lines expressing MYCN genetic-locus-based constructs demonstrate enhanced growth and oncogenic signaling compared to those solely expressing MYCN protein-coding sequences. Intriguingly, cells expressing a MYCN construct devoid of protein-coding regions but retaining noncoding elements exhibit similar oncogenic phenotypes, underscoring the significance of noncoding RNA elements in cancer development ***(Figure 3)***. These results challenge conventional paradigms and highlight the need for comprehensive investigations into the regulatory functions of noncoding RNAs in NB pathogenesis.

Further exploration of MYCN’s noncoding RNA landscape holds promise for uncovering novel mechanisms underlying NB oncogenesis and identifying potential therapeutic targets. Integrating genomic, transcriptomic, and functional studies will be instrumental in deciphering the intricate networks orchestrated by MYCN noncoding RNAs in NB cells. Additionally, efforts to elucidate the molecular mechanisms associated with MYCN mRNA in NB through advanced sequencing and copy number variation analyses present exciting avenues for future research. By embracing a holistic approach that considers both protein-coding and noncoding elements of MYCN mRNA, we can gain deeper insights into NB pathophysiology and pave the way for more effective therapeutic interventions.

## Conclusion

Neuroblastoma (NB), a challenging pediatric cancer, is often driven by the amplification of the MYCN gene. Recent research has shed light on the crucial involvement of noncoding elements within MYCN mRNA in NB oncogenesis, adding complexity to its pathology. While MYCN’s protein-coding role has been extensively studied, the regulatory functions of its untranslated regions (UTRs) and introns have been overlooked in many existing models. However, integrating full-length mRNA expression in genetic models offers a more accurate representation, aiding therapeutic development. Studies involving in vitro and in vivo models have revealed that MYCN mRNA, containing both protein-coding and noncoding elements, significantly contributes to oncogenic signaling and tumor development in NB. Notably, cells expressing MYCN mRNA devoid of protein-coding regions but retaining noncoding elements exhibit similar oncogenic phenotypes, emphasizing the importance of noncoding RNA elements in cancer development. Further exploration of MYCN’s noncoding RNA landscape holds promise for uncovering novel mechanisms underlying NB oncogenesis and identifying potential therapeutic targets. Integrating genomic, transcriptomic, and functional studies will be crucial in deciphering the intricate networks orchestrated by MYCN noncoding RNAs in NB cells. Efforts to elucidate the molecular mechanisms associated with MYCN mRNA in NB through advanced sequencing and copy number variation analyses present exciting avenues for future research. By embracing a holistic approach that considers both protein-coding and noncoding elements of MYCN mRNA, deeper insights into NB pathophysiology can be gained, ultimately leading to more effective therapeutic interventions and improved outcomes for patients battling this devastating disease.

## Methods

### Cell Culture

The neuroblastoma cell lines Neuro2a (ATCC-CCL131), H-SY-5Y (ATCC CRL-2266), and M-MSV-BALA/3T3 (ATCC CCL-163.2) were acquired from ATCC for research purposes only and were cultured in RPMI-1640 media supplemented with 10% heat-inactivated fetal bovine serum (HI-FBS). The cells were carefully grown at 37°C in a water-saturated atmosphere of 95% air and 5% CO2. All cell lines were acquired or purchased for this study and are not among the commonly misidentified ones. They were rigorously tested and found negative for mycoplasma contamination.

### Plasmids

The GFP: pPB[Exp]-EF1A>EGFP(ns):**P2A**:{mCRISPRed}, *MYCN ORF* pPB[Exp]-EF1A>EGFP(ns):**P2A**:hMYCN[NM_001293228.2]; *MYCN GL*: pPB[Exp]-EF1A>{MYCN Genetic Locus}:BGH pA-hPGK>EGFP and *Null-GL* pPB[Exp]-EF1A>{mEGFP/mycn genetic locus} were purchased from Vector Builder. Neruo2a (N2a), M-MSV BALB/3T3 (3T3) and SHSY-5Y cells were reverse-stably transfected with the above-listed plasmid, using lipofectamine™ 3000 transfection reagent (Invitrogen™) and a transposase plasmid (pRP[Exp]-mCherry-CAG>hyPBase) at a one μg concentration into six-well plates. Then, fluorescence-activated cell sorting (FACS) techniques were used to isolate transfected cell populations and use them for experiments.

### Cell Growth Assay

The cell growth assay was conducted systematically. N2a, 3T3, and SHSY-5Y cells expressing *GFP, MYCN ORF, MYCN GL, and Null GL* were used to measure proliferation rates. N2a(5*10^3cells/well), 3T3(5*10^3cells/well), and SHSY-5Y (5× 103cells / well) were seeded in 96-well plates with 10% and 2.5 % serum media for 24hr, 48hr, and 72hr. The proliferation rate was measured using the WST 8-cell proliferation assay kit (Cayman Chemical) at 450nm absorbance. The proliferation rates (n=3) were calculated based on the differences between these two-time points, providing a clear measure of cell growth.

### Western blotting

Western blots were performed with antibodies against B-actin (Cell Signaling #4970), MYCN (Santa Cruz Biotechnology #sc-53993), and GPF (Cell Signaling #2956)

### RT-qPCR

Total RNA was isolated from cells using Trizol© reagent (Life Technologies). For mRNA analysis, cDNA was prepared from 1ug RNA using a Varso kit (Life Technologies) and random hexamers. 20ng of cDNA was then used for qPCR with the IDT Primer Probe Master Mix. For mRNA expression analysis, relative copies per pg mRNA were determined using the standard curve method.

### Xenografts

N2a and 3T3 cells were transduced with GFP, MYCN ORF, MYCN GL, *and Null GL*, and 2E6 and 1E6 transduced cells were injected subcutaneously into Male StrainA and Balb/c mice, respectively. 10d-14d post-injection, mice were sacrificed, and tumors were removed and weighed. This procedure is approved by the University of Texas At Austin Institutional Animal Care and Use Committee.

